# Bacterial Conjugation in the Ruminant Pathogen *Mycoplasma agalactiae* is Influenced by Eukaryotic Host Factors

**DOI:** 10.1101/2025.02.14.638229

**Authors:** M’hamed Derriche, Laurent Xavier Nouvel, Maria Gaudino, Eveline Sagné, Elisa Simon, Hortensia Robert, Gwendoline Pot, Gilles Meyer, Christian de la Fe, Yonathan Arfi, Renaud Maillard, Christine Citti, Eric Baranowski

**Affiliations:** IHAP, Université de Toulouse, INRAE, ENVT, Toulouse, France; Ruminant Health Research Group, Departamento de Sanidad Animal, Facultad de Veterinaria, Universidad de Murcia, Murcia, Spain; Univ. Bordeaux, INRAE, UMR BFP, F-33882, Villenave d’Ornon, France

**Keywords:** bacterial conjugation, mycoplasma, mobile elements, integrative and conjugative elements, chromosomal transfer, epithelial cells, organotypic cultures

## Abstract

Bacterial conjugation plays a pivotal role in the evolution and adaptation of genome-reduced mycoplasmas. Despite their fast evolution rate, the conjugative properties of these organisms remain largely understudied, particularly *in vivo*. In the present study, the ruminant pathogen *Mycoplasma agalactiae* was used as a model organism to document the conjugative properties of mycoplasmas in environments of increasing complexity, from axenic to cell and organotypic culture conditions. Compared to axenic mating conditions, mycoplasma co-cultivation with goat epithelial cells or bovine precision-cut lung slices (PCLS) resulted in enhanced mating frequencies with high rates of *M. agalactiae* Integrative and Conjugative Element (ICEA) self-dissemination. These results were conditioned by the presence of eukaryotic cells in the culture and influenced by competition between mating partners but were not limited to *M. agalactiae*, as similar results were observed with *Mycoplasma bovis.* Mycoplasma conjugation *ex vivo* was further characterized by analyzing mycoplasma chromosomal transfer (MCT), a newly discovered mechanism of horizontal exchange of chromosomal DNA that generates mosaic genomes. Although closely associated with ICEA transfer, MCT was detected at low rates under cell and organotypic culture conditions suggesting a complex interplay between these two conjugative processes or a poor viability of the MCT progeny. Finally, mating experiments under nutrient-deprived conditions identified nucleotide stress as a potential factor influencing the modulation of mycoplasma conjugation by eukaryotic host cells. In conclusion, these results suggest that horizontal gene transfer *in vivo* is likely underestimated and provide valuable models to further studying mycoplasma conjugation *ex vivo*.

**IMPORTANCE:** Conjugation is an evolutionary shortcut that bacteria use to exchange genetic information with their neighbors. Despite the fast evolution rate of the genome-reduced mycoplasmas, their conjugative properties remain largely understudied, particularly *in vivo*. Here we used the ruminant pathogen *Mycoplasma agalactiae* to study how mycoplasmas conjugate in co-culture with hosts-derived cells and tissues. Interestingly, conjugation was stimulated when mycoplasmas were co-cultured with eukaryotic cells. This was documented by monitoring the self-propagation of a mobile genetic element known as Integrative and Conjugative Element (ICE) and the exchange of chromosomal DNA leading to the formation of mosaic genomes. While ICE transfer was observed at high frequency, only a few mosaic genomes were detected in the presence of eukaryotic cells. Further data point towards nucleotide stress as a possible factor modulating mycoplasma conjugation in cellular environments. These results suggest that mycoplasma-host interactions may stimulate conjugation *in vivo*.

## INTRODUCTION

Bacterial conjugation is the predominant mechanism of horizontal gene transfer (HGT) in microbial communities. By enabling DNA to be transferred between two bacterial cells through direct contact, conjugation facilitates the acquisition and dissemination of new adaptive traits, including antimicrobial resistance (1–3). This horizontal process is classically mediated by plasmids or Integrative and Conjugative Elements (ICE), which encode the conjugative machinery that promotes mating pair formation between donor and recipient cells and DNA transfer across a mating channel (4–7). Conjugation have been documented in a broad number of bacterial species, including mycoplasmas (8).

Mycoplasmas are fast evolving, wall-less bacteria of the class *Mollicutes*, which are restricted to an obligate parasitic lifestyle owing to the reduced genomes (9–11). Members of the genus *Mycoplasma* have a predilection for mucosal surfaces of the respiratory and genital tracts of their hosts, with some species causing chronic, often debilitating, infections in humans and animals (9, 10, 12). The control of pathogenic species is challenging, in particular due to alarming rates of antimicrobial resistance (10, 13, 14). Recently, a new mechanism of HGT known as Mycoplasma Chromosomal Transfer (MCT) has been characterized in *Mycoplasma agalactiae*, a pathogenic ruminant mycoplasma species (8, 15). MCT is an unconventional process that generates highly heterogeneous populations of mosaic genomes through the massive exchange of chromosomal DNA between mating partners (15–17). During MCT, DNA fragments originating from any part of the donor genome replace regions of a few nucleotides to several dozen kilobase pairs of the recipient chromosome at homologous sites (8, 15–17). Mainly documented *in vitro,* cumulative evidence suggests that MCT may also occurs *in vivo* (18, 19).

MCT is tightly coupled to conjugation which is mediated by a self-transmissible ICE (8, 15, 20). Widely distributed across *Mollicutes*, mycoplasma ICEs are key mediators of horizontal gene flow in these organisms (8, 21). The best characterized of these self-transmissible elements is the functional ICE of *M. agalactiae* (ICEA) (20, 22). Unlike conventional ICEs, which rely on site-specific tyrosine recombinases for their mobility, ICEA movements are mediated by a DDE transposase responsible for random, and often multiple, chromosomal integration events that generate a diverse population of transconjugants (20).

The *in vitro* study of HGT in mycoplasmas have been facilitated by the establishment of optimal mating conditions for ruminant mycoplasmas, including the use of complex axenic growth media to meet the nutritional requirements of these fastidious organisms (20, 23). However, functional genomics studies have shown that the biology of these organisms can be significantly influenced by their replicative environment, particularly when co-cultured with eukaryotic host cells (24–26). This raises questions about the influence of mycoplasma-host interactions on ICE-mediated conjugation and MCT. To address this issue, mating experiments were carried out with the model species *M. agalactiae* under replicative environments of increasing complexity aiming to mimic more closely the *in vivo* conditions. When compared to optimal axenic conditions, mating experiments revealed enhanced conjugative activity upon mycoplasma co-incubation with eukaryotic host cells or precision-cut lung samples (PCLS) with important differences among strains. The dependence of mycoplasmas on host cells for nutrient acquisition prompted us to investigate the influence of nutritional stress on the mating process. Mating experiments under nutrient-deprived conditions identified nucleotides as a potential factor influencing mycoplasma conjugation under cell culture conditions.

## RESULTS

### Mycoplasma co-cultivation with epithelial cells induces high-frequency ICEA transfer

To date, the conjugative properties of *M. agalactiae* have only been studied in axenic conditions (8). Under these conditions, ICE donor and recipient cells are co-incubated in SP4 medium after centrifugation to promote contact between the mating partners (20, 23). Since these settings differ greatly from natural growth conditions, co-cultivation of *M. agalactiae* with host epithelial cells was used to assess mycoplasma conjugative capacity *in vivo*. Mating experiments were conducted with strain PG2 and antibiotic resistance markers (Ab-tags) were used to monitor ICEA transfer from donor to recipient cells (Fig. 1, mating A). The PG2^T^[ICEA]^G^ clone was used as ICEA donor (Table 1). Its genome is characterized by the presence of (i) a functional ICEA tagged with a gentamicin marker (G-tag) to follow ICE movements and (ii) a chromosomal tetracycline marker (T-tag) to monitor chromosomal DNA exchange between mating partners (see below). The ICEA recipient partner consisted of a pool of five puromycin (P) resistant PG2 clones (further designated pPG2^P^) that was used to minimize any negative effect resulting from the chromosomal insertion of the P-tag (Table 1).

**Figure 1.**
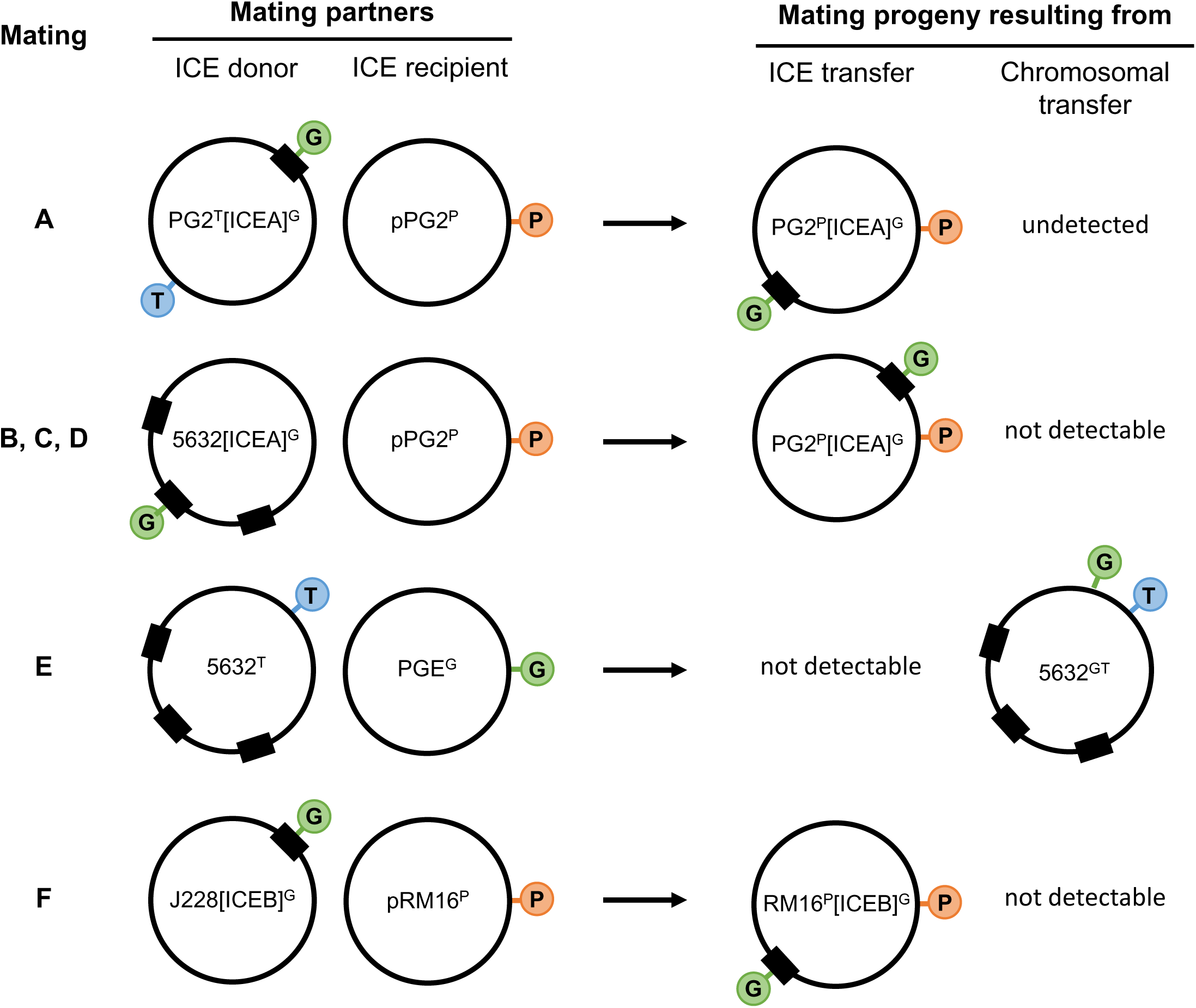
Mating experiments with *M. agalactiae* and *M. bovis*. Illustration of the different mating experiments performed in the present study. For each mating experiment (indicated with a letter), the two partners are illustrated with their specific antibiotic markers (Ab-Tag) represented by colored circles. The ICEA G-tag in green and the chromosomal T- and P-tags in blue and orange, respectively. The chromosomal ICEA is represented by a black box. The mating progenies resulting from ICEA or chromosomal transfer is illustrated, except when no progeny was detected (undetected), or when the progeny could not be detected due to the absence of an appropriate Ab-tag (not detectable).

**TABLE 1.** Ab-tagged mycoplasma strains used in mating experiments.

| Ab-tagged strains | Comments |
| --- | --- |
| ***M. agalactiae*<sup>a</sup>** |  |
| PG2<sup>G</sup> | ICEA-negative, G-tagged PG2 mutant T8.101 (24) |
| pPG2<sup>P</sup> | Pool of five ICEA-negative PG2 mutants, each having a chromosomal P-tag inserted randomly by transformation with pMT85Pur |
| PG2<sup>T</sup>[ICEA]<sup>G</sup> | ICEA-positive PG2<sup>T</sup>[ICEA *ncr19/E::mTn*]<sup>G</sup> clone 23 (22). This clone is a T-tagged PG2, which have acquired a G-tagged ICEA from 5632[ICEA]<sup>G</sup> upon mating |
| 5632<sup>T</sup> | ICEA-positive 5632 clone H3 having a chromosomal T-tag inserted outside the three ICEA regions (15) |
| 5632[ICEA]<sup>G</sup> | ICEA-positive 5632[ICEA *ncr19/E::mTn*]<sup>G</sup> clone 23 having a G-tag inserted within ICEA non-coding region 19/E (intergenic region between CDS19 and CDSE) with no influence on conjugation (22) |
| ***M. bovis*<sup>b</sup>** |  |
| pRM16<sup>P</sup> | Pool of five ICEB2-negative RM16 mutants, each having a chromosomal P2-tag inserted by transformation with pMT85Pur2 |
| J228[ICEB]<sup>G</sup> | ICEB2-positive J228[ICEB2 *ISMbov1::mTn*]<sup>G</sup> clone 277 having a G-tag inserted within ICEB2 *ISMbov1* located between ICEB2 CDS28 and 14 |

<sup>a</sup> *M. agalactiae* strains PG2 and 5632 have been previously described (19, 28). Their genome is available in databases (GenBank accession numbers CU179680.1 and FP671138.1). <sup>b</sup> *M. bovis* strains RM16 and J228 have been previously described (25, 39). Their genome sequence is available in databases (GenBank accession numbers CP077758.1 and CP068731.1).

After increasing incubation times under previously defined axenic or cell culture mating conditions, Ab-tagged partners and their dual-resistant progeny were counted by plating on selective media (Fig. 2A). Under axenic conditions, a progressive decline in mycoplasma titers of the parent strains was observed, which is likely attributable to the high initial cell density of the inoculum (10^9^ CFU/ml) classically used in this assay. Dual-resistant colonies were detected right after mixing the mating partners under axenic conditions (see 0 hours of incubation in Fig. 2A). These are likely resulting from mating events that occurred shortly after the centrifugation step used to promote contact between mycoplasma cells. In contrast, the cell culture assay was initially designed (24, 27) and conducted here with low initial density of mating partners (10^6^ CFU/ml) and resulted in bacterial growth.

**Figure 2.**
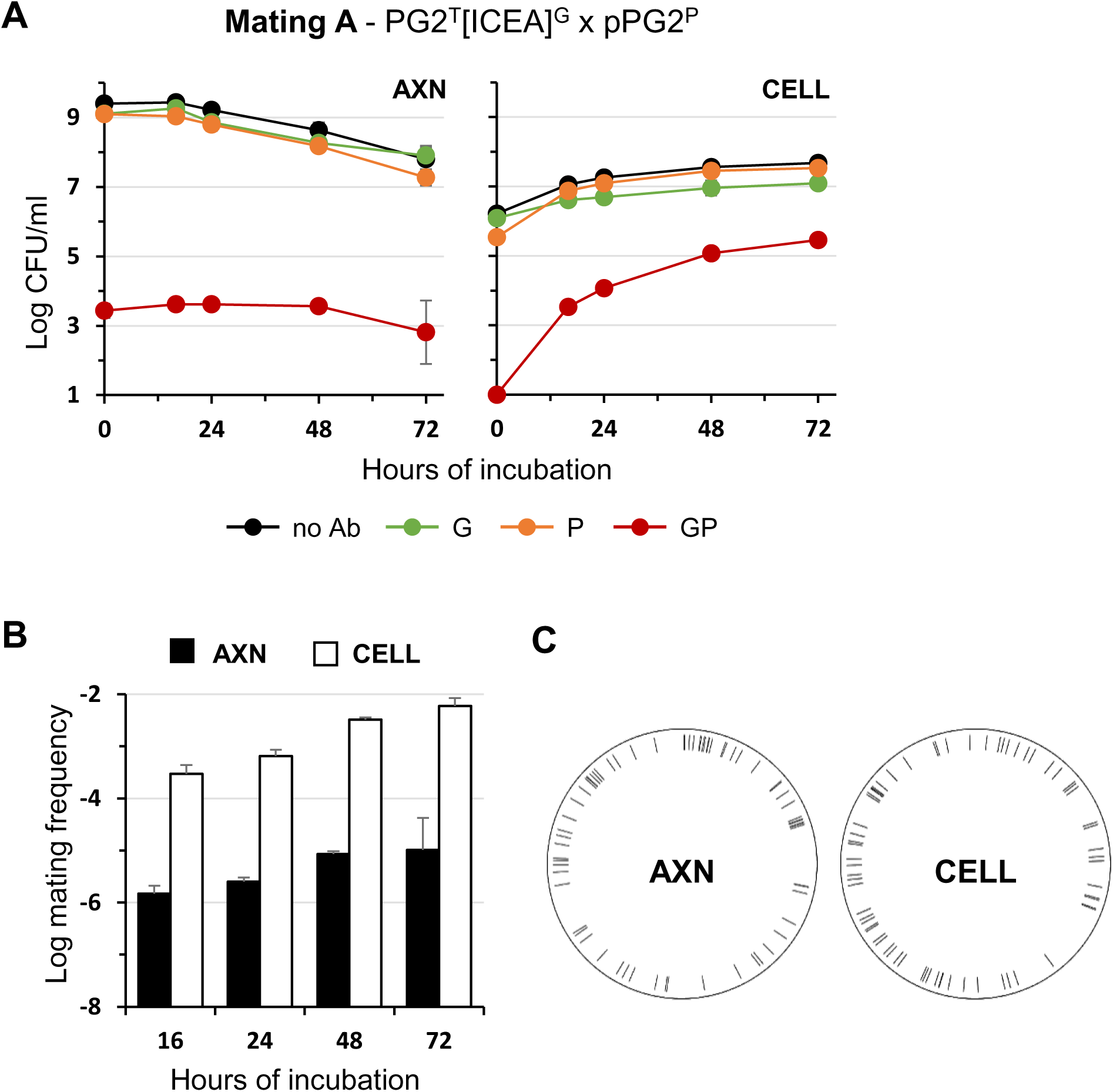
Mating experiments with *M. agalactiae* strain PG2 under axenic and cell culture conditions. (**A**) Variations in mycoplasma titers under axenic (AXN) and cell culture (CELL) mating conditions with ICEA positive and ICEA negative PG2 mating partners (Mating A in Fig. 1). Ab-tagged partners and their dual-resistant offspring were analyzed by plating on selective media. Antibiotics used for selection are color-coded (green for G; orange for P; red for G and P). Mycoplasma titers without Ab selection are shown in black. (**B**) Mating frequencies under axenic (AXN) and cell culture (CELL) conditions at different time points. Mating frequencies were calculated as the number of dual-resistant transconjugants per total CFU. (**C**) Distribution of unique chromosomal ICE integration sites identified in axenic (AXN) and cell culture (CELL) mating progeny. Data are means of at least three independent experiments. Standard deviations are indicated by error bars.

Both mating conditions generated dual-resistant colonies characterized by gentamicin and puromycin (GP) resistances (Fig. 2A) but no tetracycline and puromycin (TP) dual-resistant mycoplasmas were recovered (data not shown). ICEA transfer is therefore the main conjugative process observed between the PG2 mating partners used in these assays, since chromosomal exchange that would have conferred TP dual-resistance was not detected regardless of the conditions (Fig 1, mating A).

In each assay, the growth curve of the dual-resistant colonies mirrored that of the parents: in axenic conditions, GP dual-resistance remained stable and low (ca. 10^3^ CFU/ml) while a remarkable increase in the number of dual-resistant colonies was observed upon co-incubation with epithelial cells, reaching up to 10^5^ CFU/ml at 72 hours. Remarkably, when mating frequencies were calculated at 16 hours, the ratio of dual-resistant to total CFUs was approximately 200-fold higher in cell culture compared to axenic conditions and remained nearly stable over time (Fig. 2B and Table 2). These data suggest that the conjugative properties of *M. agalactiae* may be influenced by environmental factors such as those provided by cell culture conditions. However, mating frequencies are also likely to be influenced by differences between the two mating procedures, such as the density of mating partners, the composition of the incubation medium, or even the centrifugation step. To address these issues, axenic mating experiments were performed following the conditions set for cell culture, using low titers of mating partners and no centrifugation step (Fig. 3A). These modified axenic conditions had a negative effect on conjugation since GP dual-resistance could only be detected during the late stationary phase (48 hours) following proliferation of the mating partners (Fig 3A, SP4 medium). As expected, no or only sporadic conjugative activity was detected when the SP4 medium was replaced by DMEM medium (Fig. 3A), since mycoplasmas are unable to proliferate in cell culture media in the absence of host cells (24, 25). These results indicate that co-incubation with host cells is not only necessary for mycoplasma proliferation under cell culture conditions but also stimulates conjugation.

**Figure 3.**
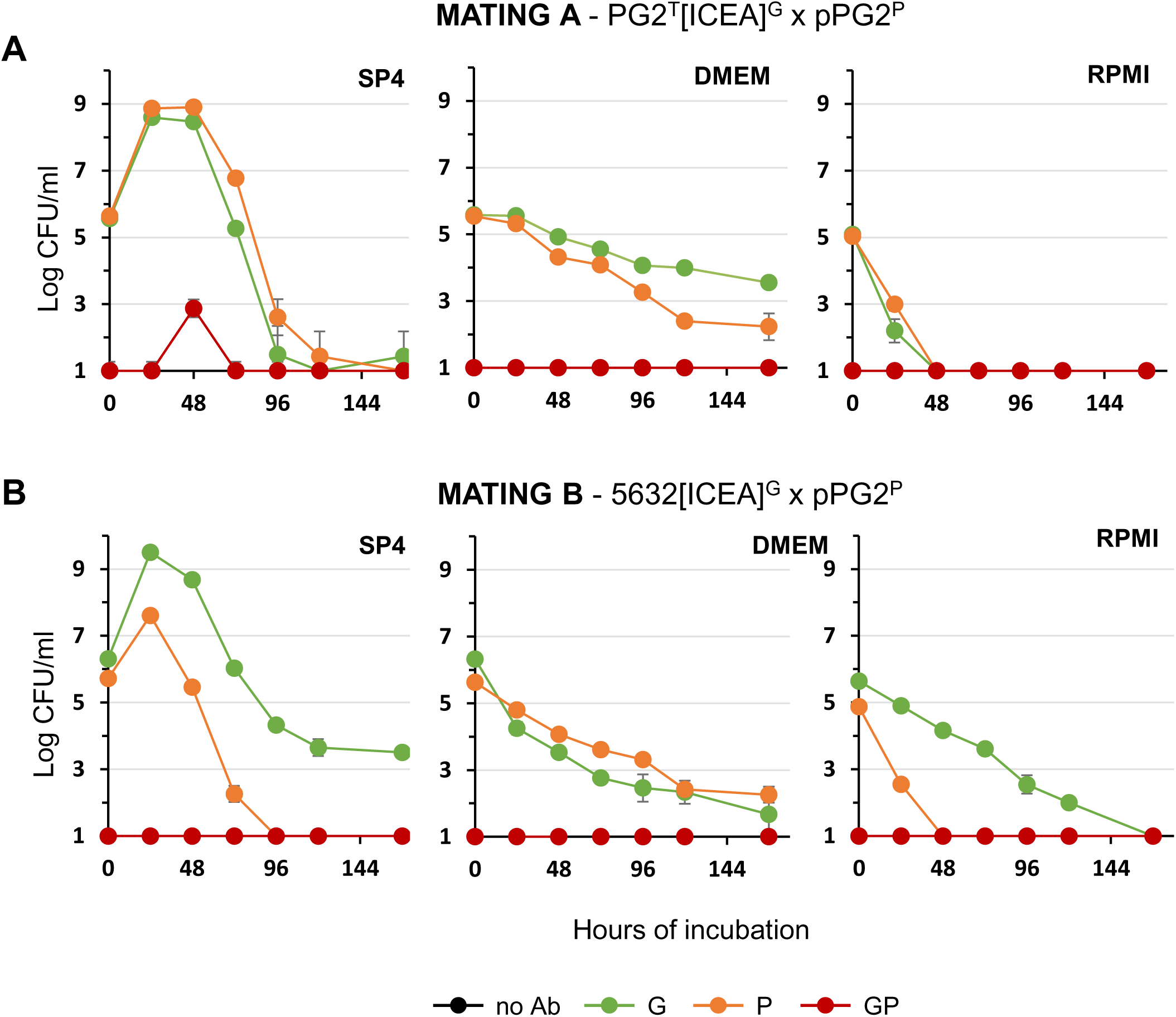
Mating experiments with *M. agalactiae* strain PG2 and 5632 in culture media. Variation of mycoplasma titers in SP4 (SP4), DMEM (DMEM), RPMI (RPMI) media upon co-incubation of PG2^T^[ICEA]^G^ and pPG2^P^ (**A**), or 5632[ICEA]^G^ and pPG2^P^ (**B**). Antibiotics used for selection are color-coded (see Fig.2). Data are the means of three independent assays. Standard deviations are indicated by error bars.

**TABLE 2.** Mating frequency of *M. agalactiae* and *M. bovis ex vivo*.

| Mating | ICE-positive partner | ICE-negative partner | Mating frequency ($\times 10^{-6}$)<sup>a</sup> | | |
| --- | --- | --- | --- | --- | --- |
|  |  |  | AXN | CELL | PCLS |
| A | PG2<sup>T</sup>[ICEA]<sup>G</sup> | pPG2<sup>P</sup> | 2.6 ± 0.4 | 670 ± 180 | 66.9 ± 96 |
| B | 5632[ICEA]<sup>G</sup> | pPG2<sup>P</sup> | 0.3 ± 0.1 | 68 ± 8.1* | 3.9 ± 3.0 |
| C | P3/CELL | pPG2<sup>P</sup> | ND | 2.4 ± 1.1 | ND |
| D | P3/AXN | pPG2<sup>P</sup> | ND | 25 ± 4.1 | ND |
| E | 5632<sup>T</sup> | PG2<sup>G</sup> | 0.3 ± 0.1 | 1.5 ± 0.7* | 0 |
| F | J228[ICEB]<sup>G</sup> | pRM16<sup>P2</sup> | 0 | 7.2 ± 4.7 | ND |

<sup>a</sup> Frequencies are given for 24 h of incubation under axenic (AXN), cell (CELL) and organotypic (PCLS)
culture conditions. (\*) Asterisks denote values obtained at 48h incubation when no mating progeny
has been selected before. ND, not determined. The values were expressed as means ± standard
deviations. The number of independent assays was 3 for each condition, except for mating C under
CELL conditions that was 9, mating B under PCLS conditions that was 13, and mating A and E under
PCLS conditions that was 16.

Axenic and cell culture mating progenies were further characterized by mapping the chromosomal ICEA-integration sites. For each mating condition, 180 dual-resistant clones were isolated at 16 hours. Individual clones were characterized by multiplex PCR amplification targeting the ICEA CDS22 and the chromosomal region surrounding the ICEA integration site in PG2^T^[ICEA]^G^ (Table S1). These amplifications (i) confirmed the presence of an ICEA in each clone and (ii) ruled out the acquisition of ICEA by chromosomal DNA exchange between mating partners (data not shown). Dual-resistant clones were then pooled, and their genomic DNA of was extracted and sequenced using Illumina technology. Reads spanning DNA junctions between ICEA and the chromosome were mapped on the PG2 reference genome as described in Materials and Methods. Sequencing data revealed a broad distribution of ICEA-integration sites over the chromosome and identified a similar number of unique integration sites for axenic (67 sites) and cell culture (69 sites) conditions (Fig. 2C and Tables S2 and S3).

Although only 40% of ICEA integration sites were detected, the data are consistent with the previously documented absence of a specific chromosomal integration site for ICEA and further illustrate the high genetic heterogeneity resulting from ICEA transfer (8, 20). More importantly, the number and distribution of ICEA-integration sites in both mating progenies also rule out the hypothesis of the early expansion of a few highly fit, dual-resistant clones whose proliferation would have been responsible for the high dual-resistant titers detected under cell culture mating conditions.

Altogether, these results show that co-culture with host cells results in higher frequency of ICEA transfer in *M. agalactiae* when compared to axenic conditions.

### *M. agalactiae* conjugation can be influenced by the fitness of the mating partners

To further investigate the conjugative properties of *M. agalactiae* under cell culture conditions, mating experiments were conducted with strain 5632, which is genetically and phenotypically distant from PG2 (28). This was performed by using the clone 5632[ICEA]^G^ as ICEA donor and pPG2^P^ as ICEA recipient (Fig. 1, mating B). The G-tagged ICEA in 5632[ICEA]^G^ is identical to PG2^T^[ICEA]^G^ (Table 1). Of note, the 5632 naturally contains 3 ICEA copies and therefore 5632[ICEA]^G^ possesses two additional untagged ICEA copies, both of which are functional and nearly identical to the G-tagged ICEA copy (15, 22). As a consequence, GP dual-resistance only monitors the self-dissemination of a single ICEA copy from 5632[ICEA]^G^ to pPG2^P^, since the movements of the untagged ICEA copies cannot be monitored. Finally, in this experiment chromosomal DNA exchanges could not be monitored. In previous study, this event was shown to only occur from PG2 to 5632 due to difference in restriction-modification systems (15, 22). Here, chromosomal DNA transfer from pPG2^P^ to 5632[ICEA]^G^ would generate non-viable transconjugants due to the presence of a Hsd-5632 restriction recognition motif in the P resistance gene used as chromosomal tag in pPG2^P^ (29) (Table 1). Under axenic conditions, 5632[ICEA]^G^ was found to outcompete pPG2^P^ ICEA recipient cells, inducing a concomitant decrease in the selection of GP dual-resistant colonies (Fig. 4A). This is likely due to the previously documented fitness advantage of 5632 over PG2 under axenic conditions (17). Another notable difference with PG2^T^[ICEA]^G^ was the lack of dual-resistant colonies selected shortly after the centrifugation step (compare 0 hours of incubation in Fig. 2A and 4A). This result suggests either a resistance of 5632[ICEA]^G^ to centrifugation-induced fusion with pPG2^P^, or a difference in ICEA activation between these two strains.

**Figure 4.**
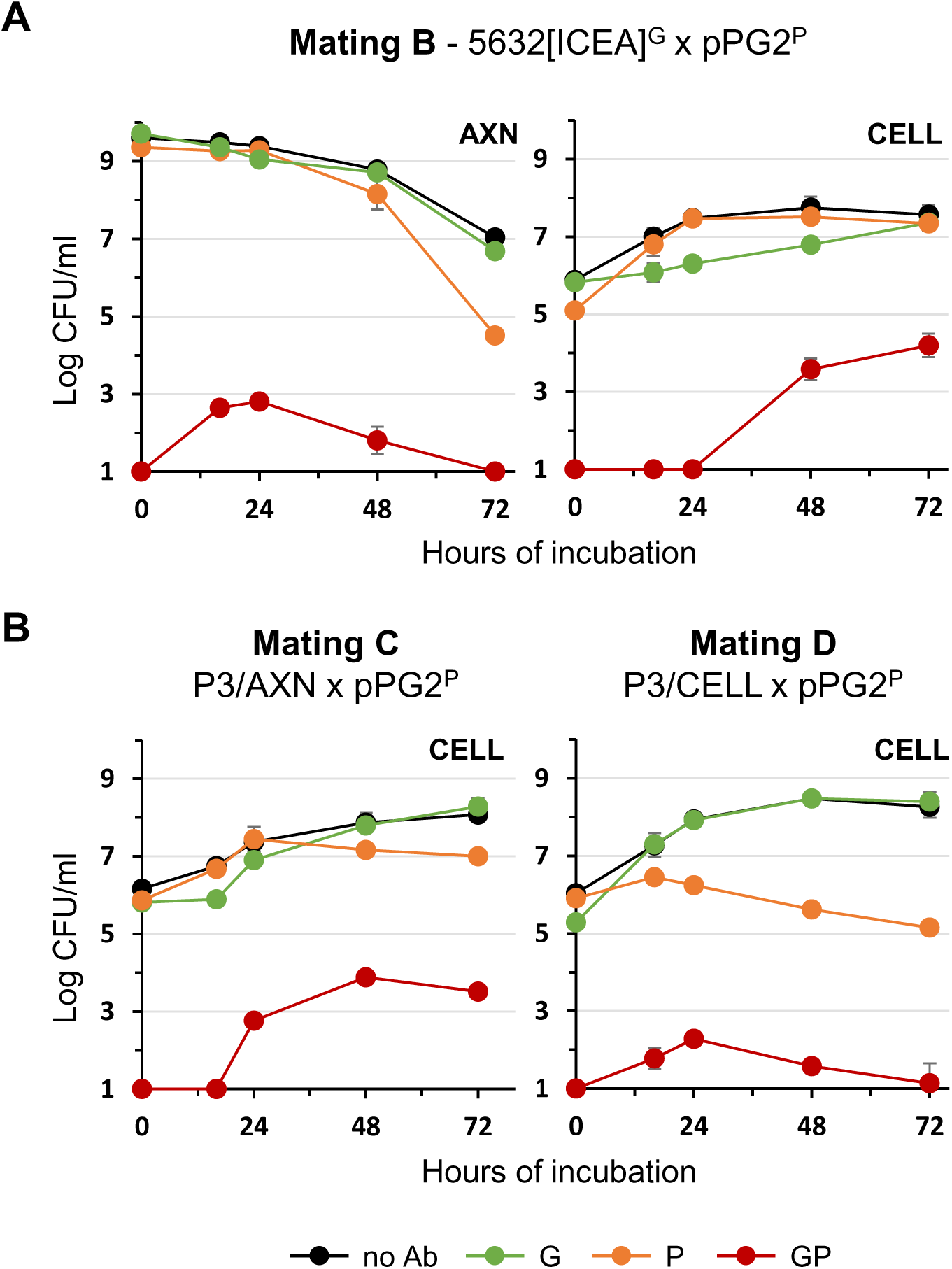
Mating experiments with *M. agalactiae* strain 5632 under axenic and cell culture conditions. (**A**) Variation of mycoplasma titers under axenic (AXN) and cell culture (CELL) mating conditions with the ICEA positive 5632 and the ICEA negative PG2 mating partners (Mating B in Fig. 1). (**B**) Conjugative properties of P3/AXN and P3/CELL in cell culture (CELL) mating conditions (Mating C and D in Fig.1). Antibiotics used for selection are color-coded (see Fig.2). Data are the means of at least three independent assays. Standard deviations are indicated by error bars.

Despite the fitness advantage of 5632 over PG2 under axenic conditions (17), 5632[ICEA]^G^ showed delayed growth under cell culture conditions, reaching titers comparable to pPG2^P^ only after 72 hours (Fig. 4A). Similarly, the detection of dual-resistant colonies was delayed by 48 hours when compared to mating with PG2^T^[ICEA]^G^. As expected, no conjugative activity could be detected under axenic condition conducted with cell culture settings, either in SP4 or DMEM medium (Fig. 3B). These results highlighted important differences in the conjugative properties of PG2^T^[ICEA]^G^ and 5632[ICEA]^G^ and identified growth competition between mating partners as an important factor affecting conjugation.

Further demonstration of how partner fitness can influence conjugation came from mating experiments with cell culture-propagated populations of 5632[ICEA]^G^ (Fig. 4B). Three populations (designated P3/CELL) were generated by three serial passages in cell culture, and a control population (designated P3/AXN) was obtained upon further propagation in SP4 axenic medium. When compared to the parental 5632[ICEA]^G^, P3/CELL populations showed an improved ability to proliferate in cell culture conditions without any delay in growth and a capacity to outcompete pPG2^P^ (Fig. 4B, mating C). Consistently, dual-resistant colonies were observed early after co-incubation of the two partners, followed by a concomitant decrease in dual-resistant colonies and ICEA recipient cells. As expected, the behavior of the control population under cell culture conditions was similar to that of the parental 5632[ICEA]^G^, but with enhanced growth after 24 hours of co-incubation (Fig. 4B, mating D).

In conclusion, *M. agalactiae* conjugation is influenced not only by the replicative environment but also by the relative fitness of each mating partner in that environment.

### Co-culture with host cells also promotes the self-dissemination of *M. bovis* ICEB

To further test the influence of host factors on mycoplasma conjugation, mating experiments were conducted with *M. bovis*, a ruminant mycoplasma species closely related to *M. agalactiae*. The clone J228[ICEB]^G^ was used as ICEB donor and a pool of five RM16^P^ clones (further designated pRM16^P^) as ICEB recipient (Table 1; Fig. 1, mating F). Under axenic conditions, J228[ICEB]^G^ was found to outcompete pRM16^P^ and no dual-resistant progeny was detected (Fig. 5). Remarkably, co-incubation of these *M. bovis* partners under cell culture mating conditions led to progressive accumulation of dual-resistant colonies with titers reaching nearly 10^3^ CFU/ml, despite no obvious proliferation of the mating partners (Fig. 5). These data suggest that the effect of mycoplasma co-culture with host cells on conjugation is not limited to *M. agalactiae* and can also be observed with other ruminant mycoplasma species.

**Figure 5.**
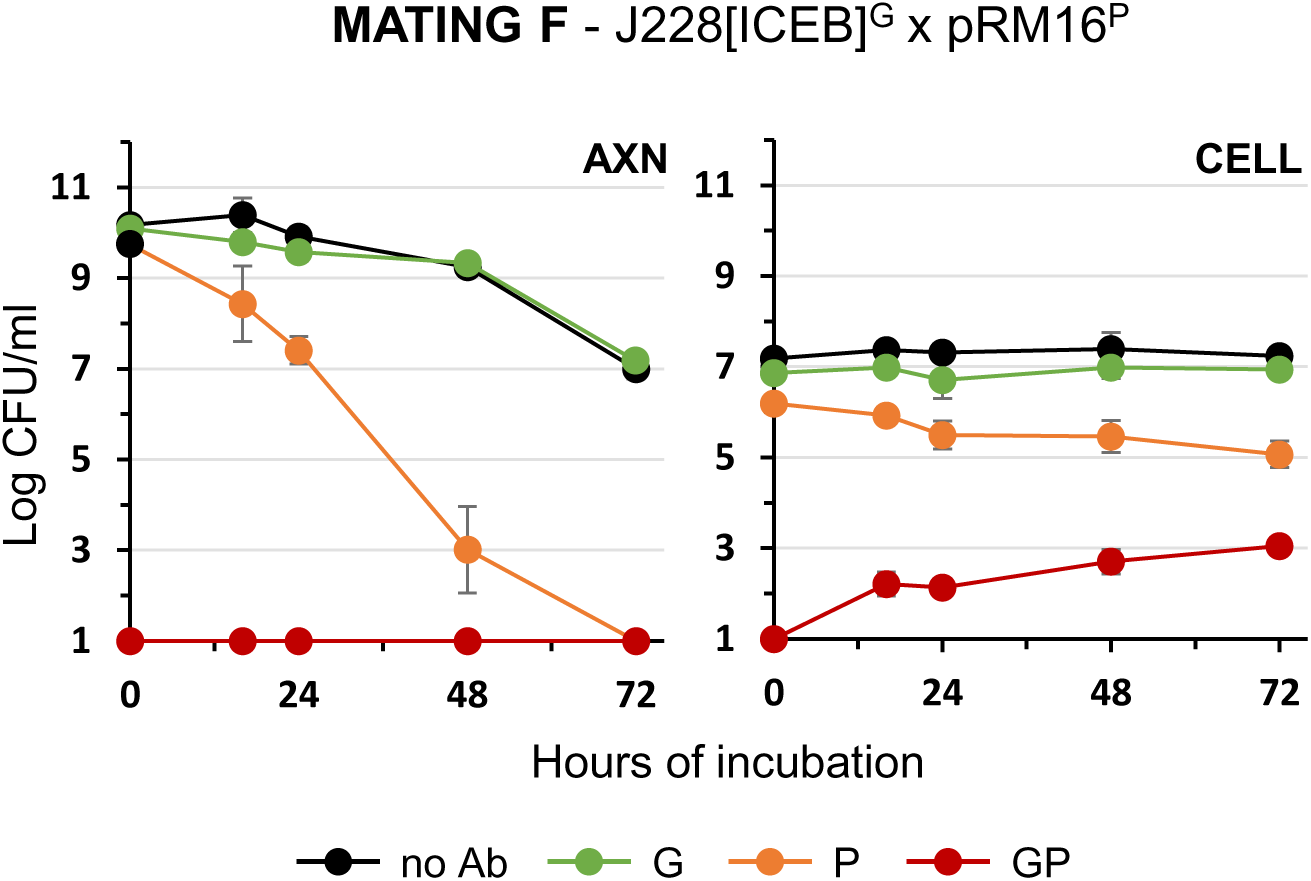
Mating experiments with *M. bovis* strain J228 under axenic and cell culture conditions. Variations in mycoplasma titers under axenic (AXN) and cell culture (CELL) mating conditions with the ICEB positive J228 and the ICEB negative RM16 mating partners (Mating F in Fig. 1). Antibiotics used for selection are color-coded (see Fig. 2). Data are means of at least three independent experiments. Standard deviations are indicated by error bars.

### Mycoplasma co-cultivation with bovine PCLS results in high-frequency ICEA transfer

Bovine precision cut lung slices (PCLS) were used to test ICEA transfer under organotypic culture conditions. PCLS were prepared from three calf lung donors and remained viable over an extended incubation period with up to 85% of the cells still viable on day 5 (see Materials and Methods). Mating experiments were conducted by using either PG2^T^[ICEA]^G^ or 5632[ICEA]^G^ as ICEA donor and pPG2^P^ as ICEA recipient (Fig. 1, mating A and B). As observed in cell culture, PG2^T^[ICEA]^G^ growth was stimulated by co-incubation with bovine PCLS (Fig. 6A). The behavior of 5632[ICEA]^G^ was dependent on the lung donor, showing either proliferation similar to PG2^T^[ICEA]^G^ with one PCLS (Fig. 6B, PCLS 1) and delayed growth with the other two (Fig. 6B, PCLS 2 and 3). Mycoplasmas remained viable even after a prolonged period of co-incubation with titers ranging from 10^7^ to 10^8^ CFU/ml. Remarkably, GP dual-resistant colonies were observed within 24 hours of co-incubation, not only with PG2^T^[ICEA]^G^, but also with 5632[ICEA]^G^. Maximum titers of dual-resistant colonies were observed after 48 to 72 hours of co-incubation and remained nearly stable even after prolonged co-incubation, likely due to limited competition between mating partners (Fig. 6A and 6B). As expected, mycoplasmas were unable to proliferate upon incubation in RPMI medium without PCLS and only sporadic conjugative activity was detected (Fig. 3A and 3B). These results indicate that ICEA transfer can occur at high frequency under organotypic culture conditions.

**Figure 6.**
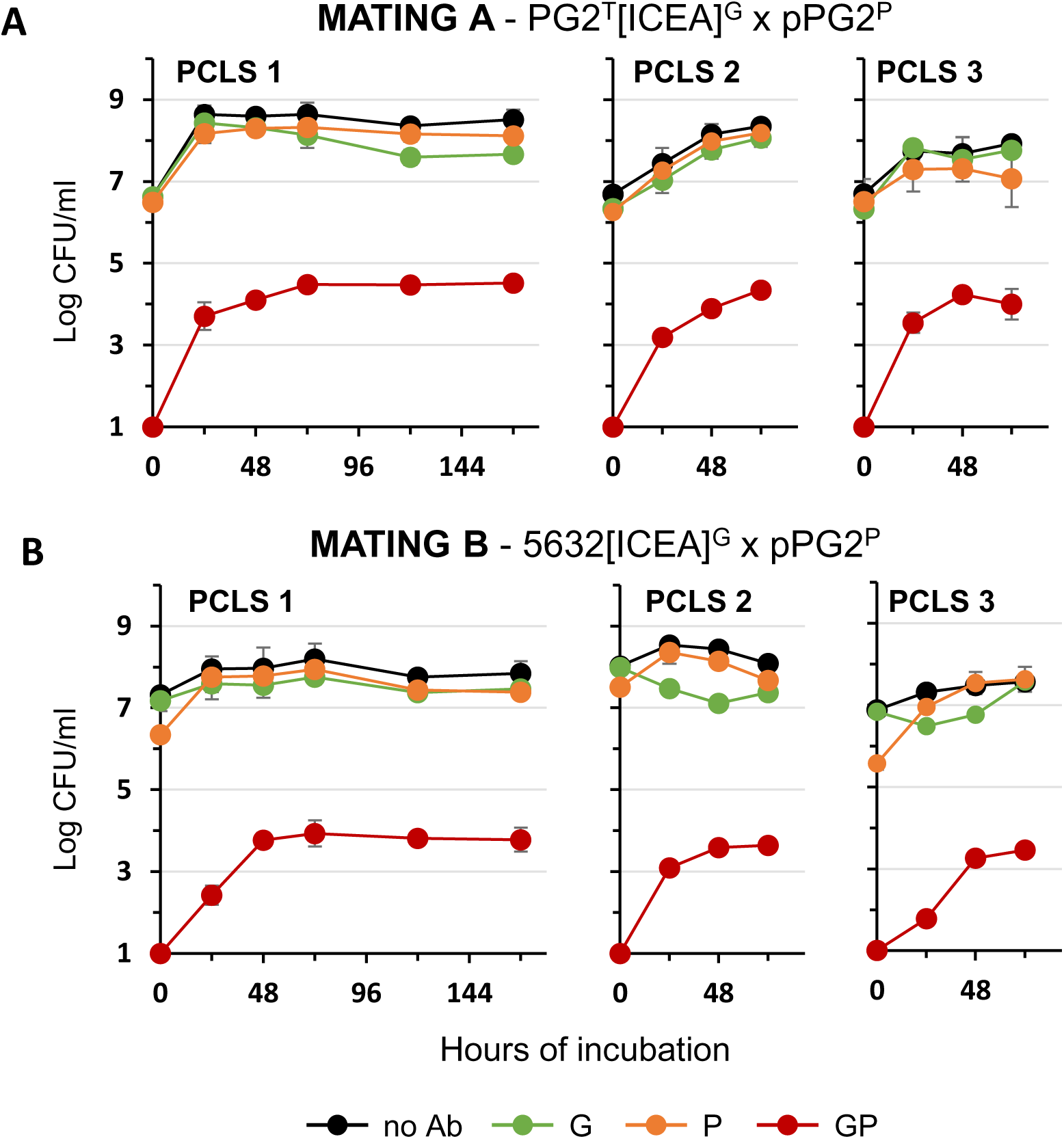
Mating experiments with *M. agalactiae* strain PG2 and 5632 under organotypic mating conditions. (**A**) Variation of mycoplasma titers under organotypic (PCLS) mating conditions with ICEA positive and ICEA negative PG2 mating partners (Mating A in Fig. 1). PCLS were from three calf donors (PCLS 1 to 3). Data from PCLS calf donor 1, 2 and 3 are the means of 6, 4 and 6 independent assays, respectively. (**B**) Variation of mycoplasma titers under organotypic (PCLS) mating conditions with the ICEA positive 5632 and the ICEA negative PG2 mating partners (Mating B in Fig. 1). Antibiotics used for the selection are indicated with a color code (see Fig. 1). Data from PCLS calf donor 1, 2 and 3 are the means of 6, 4 and 3 independent assays, respectively. Standard deviations are indicated by error bars.

### Mosaic mycoplasma genomes are generated under cell and organotypic mating conditions

The high frequency of GP dual-resistance generated upon mating with 5632[ICEA]^G^ and pPG2^P^ under cell and organotypic culture conditions led us to evaluate the rate of chromosomal transfer between these two mating partners. This was done by using a 5632^T^ ICEA positive partner and a PG2^G^ ICEA negative partner, each labeled by a specific Ab-tagged inserted in their genome (Table 1). Since the 5632^T^ ICEA positive partner has no tagged ICEA (Fig. 1, mating E), TG dual-resistant colonies will only result from the exchange of chromosomal DNA from PG2^G^ to 5632^T^ and the creation of mosaic genomes. As expected, the growth of 5632^T^ and PG2^G^ was very similar to that described for 5632[ICEA]^G^ and pPG2^P^ under both axenic and cell culture mating conditions (Fig. 7A). However, only a limited number of dual-resistant mosaic genomes could be selected in cell culture when compared to axenic mating conditions. This situation was even more dramatic upon co-incubation with bovine PCLS, with no or only a few dual-resistant mosaic genomes detected (Fig. 7B). This suggests either (i) a particular feature of the partners used in mating E, (i) a lower frequency of MCT when compared to ICEA transfer, or (iii) the formation of mosaic genomes with limited survival under cell and organotypic culture conditions. However, further studies are needed to fully understand the particular growth and mating properties of strain 5632.

**Figure 7.**
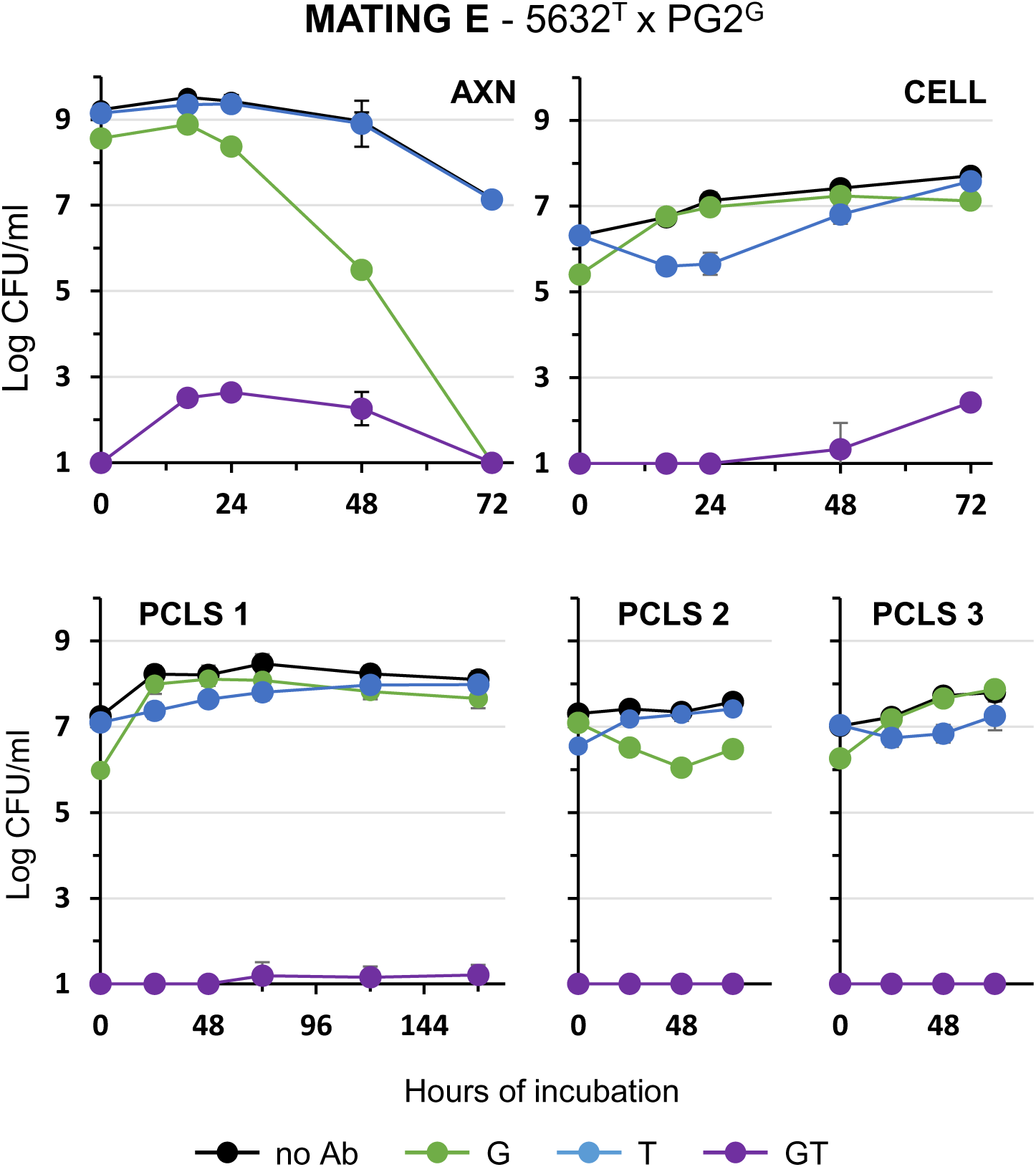
Chromosomal transfer in *M. agalactiae* under axenic, cell and organotypic culture conditions. Variation of mycoplasma titers under axenic (AXN), cell culture (CELL) and organotypic (PCLS) mating conditions with the ICEA positive 5632 and the ICEA negative PG2 mating partners (Mating E in Fig. 1). PCLS were from three calf donors (PCLS 1 to 3). Antibiotics used for selection are color-coded (blue for T; green for G; purple for T and G). Mycoplasma titers without Ab selection are shown in black. Data are the means of 3 independent assays for axenic and cell culture mating conditions. Data from PCLS 1, 2 and 3 are the means of 6, 4 and 3 independent assays, respectively. Standard deviations are indicated by error bars.

### Nucleotide stress may stimulate ICEA self-dissemination

Due to their limited metabolic capacity, mycoplasmas are dependent on their hosts for many nutrients. Under cell culture conditions, nutrients that are essential for *M. agalactiae* and *M. bovis* are provided by eukaryotic cells (24, 25, 27). Interestingly, supplementation of cell culture media with DNA can overcome this limitation and stimulate mycoplasma growth even in the absence of eukaryotic cells (25). This prompted us to study the influence of nucleotides on ICEA transfer. Mating experiments were conducted in cell culture medium supplemented with dNTPs but without eukaryotic cells. PG2^T^[ICEA]^G^ and pPG2^P^ mating partners were inoculated at low density (10^6^ CFU/ml) and analyzed by titration after 24 hours of incubation with increasing concentrations of dNTPs (Fig. 8). The addition of dNTPs was associated with a 10-fold increase in mycoplasma titer and the selection of dual-resistant colonies. Remarkably, elevated concentrations of dNTPs (ranging from 180 to 260 µM) resulted in a decrease in mating progeny with no discernible effect on mycoplasma growth, a situation that negatively affected mating frequencies. Further studies are needed to fully understand the influence of dNTPs concentration on conjugation, but these results point to nucleotide stress as a possible nutritional factor capable of stimulating ICEA transfer in mycoplasmas.

**Figure 8.**
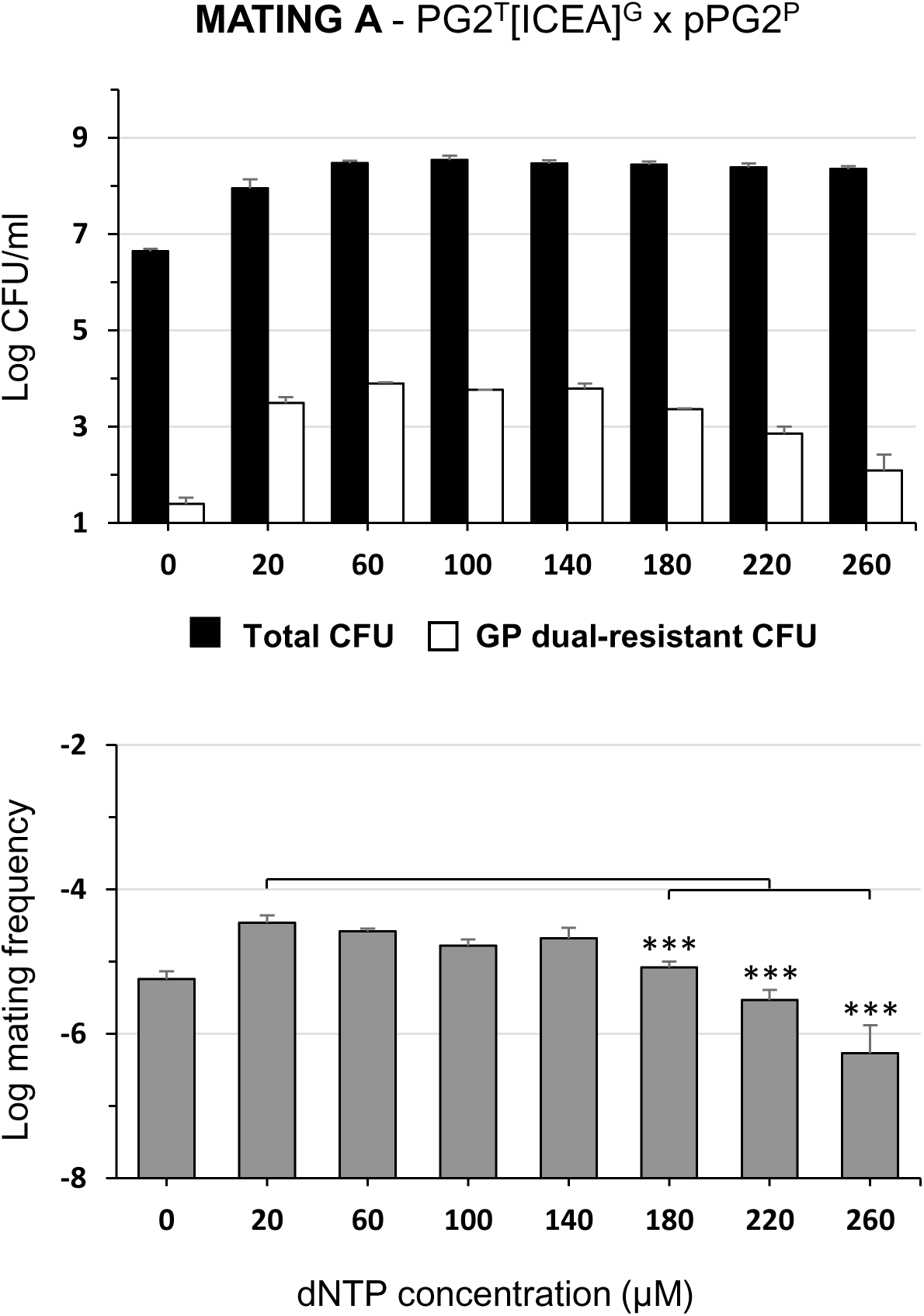
Influence of dNTPs on mating properties of *M. agalactiae* in cell culture medium. Mycoplasma titers and mating frequencies upon 24 hours co-incubation of ICEA positive and ICEA negative PG2 mating partners in DMEM medium supplemented with increasing concentration of dNTPs. No eukaryotic cells were added to the culture medium. The black and white bars show total mycoplasma titers and dual-resistant progeny selected with a combination of G and P, respectively. Data are the means of at least 3 independent assays. Standard deviations are indicated by error bars. The *p*-values were determined using two-tailed independent sample t-tests and comparing the mating frequency at a given dNTPs concentration to that at 20 µM dNTPs, which is the lower dNTPs concentration to achieve maximum mycoplasma titers without antibiotic selection (***, *p* < 0.001).

## DISCUSSION

Despite the control that mobile genetic elements exert over their own mobilization, it is becoming clear that HGT among host-associated microbes can be modulated by *in vivo* environmental conditions (1, 30–32). By using several *ex vivo* infection models, our study shows that eukaryotic cells can considerably impact the mating frequency of ruminant mycoplasma species.

### Mycoplasma ICE transfer *ex vivo*

Mating experiments with the prototype PG2 strain of *M. agalactiae* revealed elevated rates of ICEA transfer under cell culture and organotypic conditions, with a mating frequency reaching up to 7 x 10^-4^ dual-resistant colonies per total CFUs at 24 hours. This value was 200-fold higher than those measured using the established axenic method for conjugation (20, 23). This difference was not the outcome of a fast growth of several high-fitted transconjugants upon co-incubation with host cells, since the distribution of chromosomal ICEA insertion sites were comparable in both mating populations. However, ICEA transmission rates in cell culture were likely overestimated at longer incubation times due to repeated conjugation events and proliferating progeny. Without detectable MCT events, ICEA transfer was the major conjugative process stimulated by co-incubation with eukaryotic host cells.

Co-incubation with eukaryotic cells was not only able to stimulate conjugation in the prototype *M. agalactiae* strain, but also when 5632 was used as ICEA donor. However, 5632 showed reduced conjugative activity under all conditions tested. This difference is consistent with previous studies conducted under axenic mating conditions, which reported mating frequencies of 1 to 8 x 10^-6^ and 5 to 10 x 10^-8^ for PG2 and 5632 respectively (15, 20, 22). Factors influencing ICE transfer from 5632 to PG2 include (i) the number of co-resident chromosomal ICEA copies in the donor cells since ICEA movements can only be monitored for the copy that contains an Ab-tag (22), and (ii) the relative fitness of each partner that can influence the ratio of donor and recipient cells.

Finally, co-incubation with eukaryotic cells also stimulated conjugation in *M. bovis* providing the first report of ICEB transfer in this species. Remarkably, cell culture mating conditions provide a valuable model to study conjugation with this economically important species, as no activity could be detected under axenic conditions developed for ruminant mycoplasmas (23).

### Mycoplasma chromosomal transfer *ex vivo*

Mycoplasma ICEs were found to play a central role in MCT which requires a functional ICE in one of the mating partners (8, 15, 20). In contrast to ICE transfer, MCT was only sporadically observed under cell and organotypic culture conditions when 5632 was used as ICEA donor. Further studies are needed to fully understand this observation, but the mosaic genomes resulting from MCT may have reduced viability in these two cellular environments. Another hypothesis is suggested by our recent study that identified mycoplasma restriction-modification RM systems as key in controlling chromosomal transfer (29). Indeed, RM systems may influence the polarity of DNA transfers by fragmenting the unmethylated chromosome of one partner, thereby facilitating its incorporation into the methylated chromosome of the second partner. Consequently, MCT may be influenced by the composition of the RM systems in each partner and by changes in their DNA methylation profile. This hypothesis consistent with the absence of chromosomal exchanges between two partners having the same chromosomal background, such as mating experiments with two PG2 partners that failed to produce any MCT progeny under all the conditions tested. Whether adaptation of 5632 to cell and organotypic culture conditions induces changes in DNA methylation is unknown, but it is increasingly recognized that epigenetic modifications in bacteria can respond to the environment and modulate cell cycle, gene expression, and virulence (33, 34). In the human pathogen *Mycoplasma pneumoniae*, the methylation status is subject to variation as a function of the growth phase (35). Further studies are needed to characterize the changes associated with 5632 growth *ex vivo* and to determine their potential impact on MCT.

### Nucleotides stress as a potential trigger of mycoplasma conjugation

Nucleotide metabolism is emerging as critical to *M. bovis* survival and virulence (25, 26, 36, 37) and may also be key to *M. agalactiae* interaction with host cells and HGT. Indeed, dNTPs were found (i) to overcome the dependence of *M. agalactiae* on eukaryotic cells for proliferation in cell culture, and (ii) to inhibit bacterial conjugation at high concentrations. This suggests that mycoplasma conjugation under cell culture conditions may be stimulated by a nutritional stress resulting from fluctuations in nucleotide levels. Further studies are needed to elucidate the impact of nucleotide stress on the physiology of mycoplasma cells. However, it is known that starvation can lead to DNA damage and the induction of the SOS response in bacteria, a physiological condition known to trigger the activation of several ICEs in a RecA-dependent manner (5). Nevertheless, DNA repair mechanisms in mycoplasmas are still poorly understood and no SOS response has been documented except for an SOS-like response in *Mycoplasma gallisepticum* (38).

Finally, while the dependence of *M. agalactiae* on eukaryotic cells can be overcome by dNTP supplementation, eukaryotic cells were required to achieve maximal conjugation rates. Indeed, dNTP mating frequencies remain approximately 10-fold lower than those observed under normal cell culture conditions (compare mating frequencies at 24 h in Fig. 2B and 8). This suggests that multiple factors are likely involved, such as cell surface adhesion, which may facilitate the contact needed for mycoplasma conjugation to occur.

In conclusion, this study illustrates how eukaryotic host factors can promote conjugation in mycoplasmas and suggests that *in vivo* HGT events are likely underestimated. It also provides valuable models for future studies on mycoplasma conjugation *ex vivo*.

## MATERIALS AND METHODS

### Mycoplasmas and culture conditions

Ab-tagged *M. agalactiae* and *M. bovis* strains used in this study are described in Table 1. *M. agalactiae* strains PG2 and 5632 (19, 28) differ in their ICEA content, with strain 5632 having three functional, almost identical ICEA copies, while strain PG2 contains only vestigial forms (14, 16). Similarly *M. bovis* strains RM16 and J228 (25, 39) differ in their ICEB content, with strain J228 having two chromosomal ICEB-2 copies, while strain RM16 contains no ICEB (18).

Stock cultures were produced by growing mycoplasmas at 37°C in SP4 medium (40) supplemented with 100 µg/ml ampicillin (Sigma-Aldrich) and stored at -80°C. When needed, gentamicin (50 µg/ml; Gibco), puromycin (10 µg/ml; Thermo Fisher) and tetracycline (2 µg/ml; Sigma-Aldrich), alone or in combination, were added to the medium. Spontaneous resistance in stock cultures was tested by plating on selective media. Since mycoplasma growth cannot be monitored by optical density, mycoplasma titers were determined based on colony counts on solid SP4 media after 4 to 7 days of incubation at 37°C using a binocular stereoscopic microscope (24). The detection limit for mycoplasma titration was 100 CFU/ml, except for the detection of mating progeny, which was 10 CFU/ml.

### Genetic tagging of mycoplasmas with antibiotic markers

Selective antibiotic markers were introduced randomly in the mycoplasma genome by transforming mycoplasma cells with plasmid pMT85, which carries a gentamicin (G) resistance mini-transposon (mTn) (41), or its derivatives, pMT85-Tet, pMT85-Pur and pMT85-Pur2, in which the G resistance was replaced by a tetracycline (T) or a puromycin (P) resistance (20, 24). Plasmid pMT85-Pur2 encodes a codon optimized version of the puromycin resistance *pac* gene and a mutation at the Hsd-5632 recognition motif 5′-A^m6^YC(N)5KTR-3′ (29). J228[ICEB]^G^ was selected from a library of 300 mutants generated by random transposon mutagenesis and identified by its capacity to generate dual-resistant progeny upon mating with pRM16^P^ under cell culture conditions (see below). Whole-genome sequencing of PG2^T^[ICEA]^G^ identified the integration site of the G-tagged ICEA at PG2 genomic position 703341 (GenBank accession number: CU179680.1) corresponding to the N-terminal region of a hypothetical protein with unknown function encoded by MAG6030. The PG2 chromosomal T-tag is located 83 nucleotides upstream of MAG1620 (genomic position 188668), which encode a hypothetical protein.

### Epithelial cells and organotypic cultures

The T-antigen immortalized goat milk epithelial cell (TiGMEC) line was kindly provided by C. Leroux (UMR 754 INRAE, Université Lyon 1, Lyon, France) (27, 42). Cells were grown in Dulbecco’s modified Eagle’s medium (DMEM) (high glucose, sodium pyruvate, and GlutaMAX-I; Gibco) supplemented with non-essential amino acids (NEAA; Gibco) and 10% heat-inactivated fetal bovine serum (FBS, Gibco) (24, 27). The mycoplasma-free status of the cell line was tested by using a genus-specific PCR (43).

The precision bovine lung slices (PCLS) used in this study were obtained from another study (44). Briefly, lung tissues were collected from 3 to 6-week-old male calves. Lung donors that were PCR-positive for respiratory pathogens were excluded (44). PCLS of 100 µm thickness were prepared from 8 mm punch biopsies of cranial and accessorial lobes using the Krumdieck MD6000 tissue slicer (Alabama Research & Development) as described (44). PCLSs in 24-well plates were washed with RPMI medium (Gibco) supplemented with 10% FBS and 1% penicillin-streptomycin (10,000 U/mL penicillin and 10 mg/mL streptomycin, Pan Biotech). After overnight recovery at 37°C with 5% CO_2_ and two additional washes in RPMI, mating experiments were performed in RPMI supplemented with 10% FBS, amphotericin B (2.5 μg/mL; Sigma-Aldrich) and ampicillin (0.3 mg/mL; Sigma-Aldrich). PCLS viability data were published in Gaudino *et al*., 2023 (44).

### Mycoplasma mating experiments

Mating experiments with Ab-tagged mycoplasmas were conducted under axenic, cell and organotypic culture conditions. Axenic mating conditions were as previously described (20, 23). Briefly, cultures of donor and recipient mycoplasmas were mixed in a 1:1 CFU/CFU ratio, centrifuged for 5 min at 8,000 x *g* at room temperature, resuspended in 1 ml of SP4 medium, and further incubated a 37°C. At the end of the incubation period, mycoplasmas were plated onto SP4 solid media supplemented with the appropriate antibiotics for selecting either the parental populations or the transconjugant progeny or without antibiotics for total cell count. Cell culture mating experiments were performed by inoculating donor and recipient mycoplasmas to epithelial cell cultures. TiGMEC cells were seeded in 6-well plates (Falcon) at a density of 4 x 10^4^ cells/cm² and inoculated at a mycoplasma/cell ratio of ca. 5 CFU/cell (10^6^ CFU/ml). Organotypic culture mating experiments were performed by inoculating ca. 10^6^ CFU of each mycoplasma partner to individual PCLS in RPMI medium. At different times post-inoculation, mycoplasma titers were determined by CFU titration on selective SP4 solid media following three freeze-thaw cycles of cell and organotypic cultures. Mycoplasma cell viability is not affected by limited freeze-thaw cycles due to the cryoprotective effect of the serum. For mating A, B, E and F (Table 2), ten dual-resistant colonies were controlled by PCR amplifications using primers targeting the resistance genes used for their selection (Table S3). For mating A, B and E, these dual-resistant colonies were further tested with PCR amplifications using primers targeting CDS22 in ICEA (Table S3).

### Mycoplasma whole-genome sequencing and bioinformatic mapping of ICE-integration sites

Mycoplasma genomic DNA was purified by phenol/chloroform extraction and ethanol precipitation (45). Whole-genome sequencing was performed using Illumina sequencing technology (NovaSeq PE 150). Raw sequencing data were generated at Eurofins Genomics (Germany). Bioinformatics analyses were conducted using the GenoToul Bioinformatics facility, Toulouse, France (http://bioinfo.genotoul.fr/).

To map ICE-integration sites from large pools of transconjugants, Illumina raw sequencing data, after quality control and sequencing adapter trimming, were filtered to select direct and indirect reads spanning DNA junctions between ICEA and the mycoplasma chromosome. This was done by selecting reads containing nucleotide sequences GGAACTGATATAAGAAAGTG or reverse complement CACTTTCTTATATCAGTTCC corresponding to 5’-ICEA ends (hereafter referred to as CDS1 reads) and nucleotide sequences CCCACTTAATACTTTCATTC or reverse complement GAATGAAAGTATTAAGTGGG corresponding to 3’-ICEA ends (hereafter referred to as CDS22 reads). Selected reads were mapped on the PG2 reference genome (NCBI accession number CU179680) with BWA_MEM (46). BAM files corresponding to aligned reads were processed with BEDTools Intersect v2.29.1 (47) to identify overlapping CDS1 and CDS22 reads. The intersection files, generated by BEDTools Intersect v2.29.1, were imported into Artemis 16.0.0 (48), where each CDS1/CDS22 intersection was converted into a feature on the PG2 reference genome. Features were created based on the number of reads at each potential ICEA insertion site, with a minimum number of four reads for each ICEA extremity. This cutoff was selected after manual reconstitution and inspection of each features to identify false ICEA insertion sites. Selected ICEA integration sites were further examined to confirm the presence of an 8-nucleotide direct repeat on the overlap between CDS1 and CDS22 reads, a which is a characteristic signature of ICEA integration events.

## Data Availability

The datasets for this study can be found at https://www.ebi.ac.uk/, accession numbers: ERR14088891, ERR14088864, ERR14088863.

## Acknowledgments

We thank R. Herrmann for providing plasmid pMT85 and C. Leroux for TiGMEC cells. We also thank B. Collard for excellent technical assistance.

## Author Contributions

All authors read and approved the version submitted for publication.

**M. Derriche:** Conceptualization, Data curation, Formal analysis, Investigation, Validation, Visualization, Writing - original draft.

**LX. Nouvel:** Conceptualization, Data curation, Formal analysis, Funding acquisition, Methodology, Supervision, Validation, Writing - review & editing.

**M. Gaudino:** Methodology, Resources,

**E. Sagné:** Investigation, Methodology, Resources, Supervision, Validation.

**E. Simon:** Investigation, Methodology, Validation.

**H. Robert:** Investigation, Validation.

**G. Pot:** Investigation, Methodology, Validation.

**G. Meyer:** Methodology, Resources.

**C. de la Fe:** Funding acquisition, Resources.

**Y. Arfi:** Conceptualization, Funding acquisition, Project administration, Writing - original draft, Writing - review & editing.

**R. Maillard:** Funding acquisition, Resources.

**C. Citti:** Conceptualization, Funding acquisition, Project administration, Writing - original draft, Writing - review & editing.

**E. Baranowski:** Conceptualization, Data curation, Funding acquisition, Methodology, Project administration, Resources, Supervision, Validation, Visualization, Writing - original draft, Writing - review & editing.

