## Supplemental data for "Bacterial Conjugation in the Ruminant Pathogen *Mycoplasma agalactiae* is Influenced by Eukaryotic Host Factors"

### SUPPLEMENTARY TABLES

**Table S1.** PCR amplifications and oligonucleotide primers

| PCR target | Primer name | Sequence (5’ -> 3’) |
|---|---|---|
| ICEA CDS22 | ORF22-F-ICE | TGAGACCAGCAAGCTGAAGA |
| | ORF22-R-ICE | TCTGTATCAATCTGAATTGCATCAT |
| ICEA integration site in PG2$^{T}$[ICEA]$^{G}$ | ICE MAG6030 F | AGGCTAAAAGTGCTTTGGAG |
| | ICE MAG6030 R | GCTTTGAGTGAAATTGAGAGTCA |
| Gentamicin marker | GM1 | ACATGAATTACACGAGGGC |
| | GM2 | GTTCTTCTTCTGACATAGTAG |
| Puromycin marker | PuroF | GTTGCTGTTTGGACTACTCCTG |
| | PuroR | CACCAAGTTCTAGGACCTTCAGG |
| Puromycin marker P2 | PuroF2 | GTTGCTGTTTGGACTACTCCAG |
| | PuroR | See above |
| Tetracycline marker | IntMtet1 | TGGCGTACAAGCACAAACTC |
| | IntMtet2 | GCAAAGTTCAGACGGACCTC |

**Table S2.** Chromosomal ICEA insertion sites identified in a pool of 180 transconjugants selected under axenic mating conditions

| Genomic position [a] | Orientation [b] | CDS [c] | Genomic position [a] | Orientation [b] | CDS [c] |
|---|---|---|---|---|---|
| 002717 | + | MAG0020/MAG0024 | 372071 | + | MAG3110 |
| 003253 | + | MAG0030 | 413783 | + | MAG3480 |
| 008397 | + | MAG0080 | 455122 | - | MAG3850/MAG3860 |
| 013347 | + | MAG0130/MAG0140 | 456852 | - | MAG3860 |
| 020506 | + | MAG0190/MAG0200 | 468747 | + | MAG3950 |
| 021461 | - | MAG0200 | 497746 | - | MAG4240 |
| 024628 | - | MAG0240 | 505879 | + | MAG4330 |
| 026019 | + | MAG0260 | 513601 | - | MAG4390 |
| 029739 | - | MAG0280/MAG0290 | 538623 | - | MAG4590/MAG4600 |
| 030522 | - | MAG0290 | 565501 | + | MAG4840 |
| 033736 | - | MAG0340 | 575347 | + | MAG4940 |
| 048595 | - | MAG0390 | 578412 | - | MAG4960/MAG4970 |
| 049926 | - | MAG0390 | 633991 | + | MAG5470/MAG7520 |
| 060955 | + | MAG0510 | 648596 | - | MAG5630/MAG5640 |
| 068280 | - | MAG0570/MAG7500* | 656807 | + | MAG5690 |
| 090928 | - | MAG0760 | 661640 | + | MAG5730 |
| 122520 | + | MAG1060 | 663779 | - | MAG5730/MAG5740 |
| 123514 | - | MAG1070/MAG1080 | 679911 | - | MAG5880 |
| 135864 | + | MAG1210/MAG1220 | 689360 | + | MAG5950 |
| 148703 | + | MAG1310 | 695597 | + | MAG5990 |
| 166809 | - | MAG1450 | 711743 | - | MAG6090/MAG6100 |
| 169797 | + | MAG1470 | 732272 | - | MAG6160 |
| 170568 | + | MAG1470/MAG1480 | 747994 | + | MAG6300 |
| 173360 | - | MAG1500/MAG1510 | 758996 | + | MAG6410 |
| 174562 | - | MAG1510 | 761012 | + | MAG6420 |
| 176252 | - | MAG1520 | 761768 | - | MAG6430/MAG6440 |
| 244510 | - | MAG2120 | 765983 | - | MAG6480 |
| 245246 | + | MAG2130 | 769728 | - | MAG6520 |
| 252282 | + | MAG2190 | 773049 | - | MAG6560 |
| 288386 | + | MAG2480 | 790430 | + | MAG6730 |
| 306737 | + | MAG2640 | 798250 | - | MAG6820 |
| 328311 | + | MAG2810 | 819722 | + | MAG7040/MAG7050 |
| 343884 | - | MAG2880 | 846965 | + | MAG7280/MAG7285 |
| 347535 | - | MAG2920 | | | |

[a] ICEA Insertion sites have been defined based on the sequence of *M. agalactiae* strain PG2 (GenBank accession number: CU179680.1); [b] orientation of ICEA relative to the genome; [c] mnemonic of the coding sequence (CDS) in which the ICEA is integrated; when ICEA is integrated in a non-coding region, upstream and downstream CDS are indicated; * MAG7500 is located between MAG0570 and MAG0580

**Table S3.** Chromosomal ICEA insertion sites identified in a pool of 180 transconjugants selected under cell culture mating conditions

| Genomic position [a] | Orientation [b] | CDS [c] | Genomic position [a] | Orientation [b] | CDS [c] |
|---|---|---|---|---|---|
| 012795 | + | MAG0120/MAG0130 | 500148 | + | MAG4270 |
| 027594 | - | MAG0270/MAG0280 | 529095 | + | MAG4510 |
| 031912 | + | MAG0310/MAG0320 | 534625 | + | MAG4580 |
| 037621 | + | MAG0370 | 545123 | - | MAG4650 |
| 048822 | + | MAG0390 | 549591 | + | MAG4710 |
| 060916 | - | MAG0510 | 562561 | + | MAG4810 |
| 067751 | - | MAG0570 | 566387 | + | MAG4850/MAG4860 |
| 091470 | - | MAG0760 | 580342 | + | MAG4980 |
| 102881 | - | MAG0900 | 582341 | - | MAG5000 |
| 123163 | + | MAG1070 | 590039 | - | MAG5050 |
| 125437 | - | MAG1100 | 594324 | + | MAG5080 |
| 129551 | + | MAG1160 | 601868 | - | MAG5140 |
| 173108 | - | MAG1500 | 634181 | - | MAG5470/MAG7520* |
| 176037 | - | MAG1520 | 640291 | + | MAG5570 |
| 202959 | - | MAG1720 | 645279 | + | MAG5620 |
| 211481 | + | MAG1810 | 656425 | + | MAG5690 |
| 216511 | + | MAG1830/MAG1840 | 663274 | + | MAG5730 |
| 245201 | + | MAG2130 | 680355 | - | MAG5890 |
| 258414 | + | MAG2220 | 682881 | + | MAG5900 |
| 259713 | - | MAG2286/MAG2230 | 691503 | - | MAG5960 |
| 269650 | - | MAG2320 | 703018 | + | MAG6030 |
| 270644 | - | MAG2330 | 737079 | - | MAG6210/MAG6220 |
| 348006 | + | MAG2920 | 739831 | + | MAG6230 |
| 391337 | - | MAG3280 | 745143 | - | MAG6290 |
| 399791 | + | MAG3380/MAG3390 | 745711 | - | MAG6290 |
| 406268 | + | MAG3420/MAG3430 | 746707 | + | MAG6290 |
| 435944 | - | MAG3680 | 747723 | + | MAG6290 |
| 451726 | + | MAG3810 | 755031 | - | MAG6380 |
| 468642 | + | MAG3940/MAG3950 | 771645 | - | MAG6540 |
| 478287 | + | MAG4070 | 794535 | - | MAG6780 |
| 478947 | + | MAG4080 | 830651 | - | MAG7120/MAG7130 |
| 478949 | - | MAG4080 | 832984 | + | MAG7140 |
| 483866 | - | MAG4130 | 844782 | + | MAG7270 |
| 493804 | - | MAG4210 | 871348 | + | MAG7440/MAG7450 |
| 497527 | - | MAG4230/MAG4240 | | | |

[a] ICEA Insertion sites have been defined based on the sequence of *M. agalactiae* strain PG2 (GenBank accession number: CU179680.1); [b] orientation of ICEA relative to the genome; [c] mnemonic of the coding sequence (CDS) in which the ICEA is integrated; when ICEA is integrated in a non-coding region, upstream and downstream CDS are indicated; * MAG7520 is located between MAG5470 and MAG5480.

#### SUPPLEMENTARY FIGURE

**Mating A** (PG2$^{T}$[ICEA]$^{G}$ x pPG2$^{P}$)

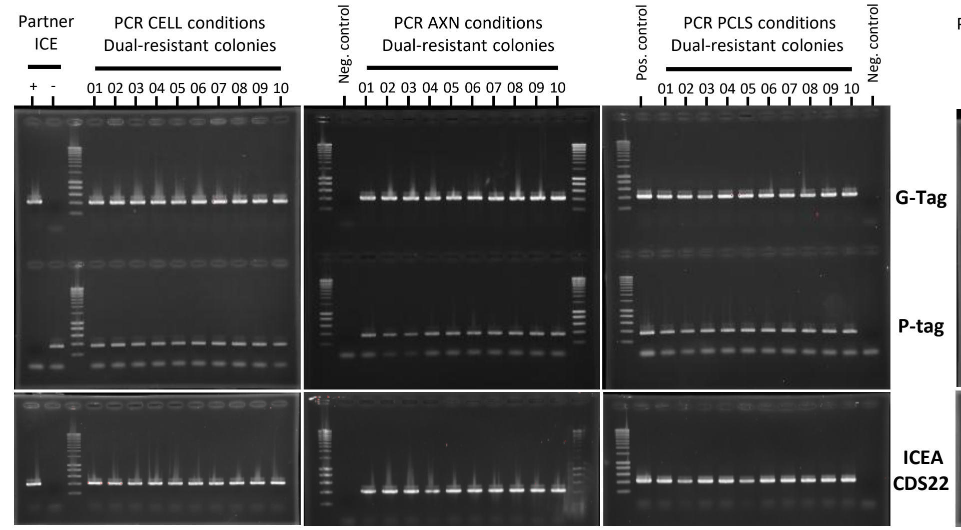


**Mating B** (5632[ICEA]G x pPG2P)

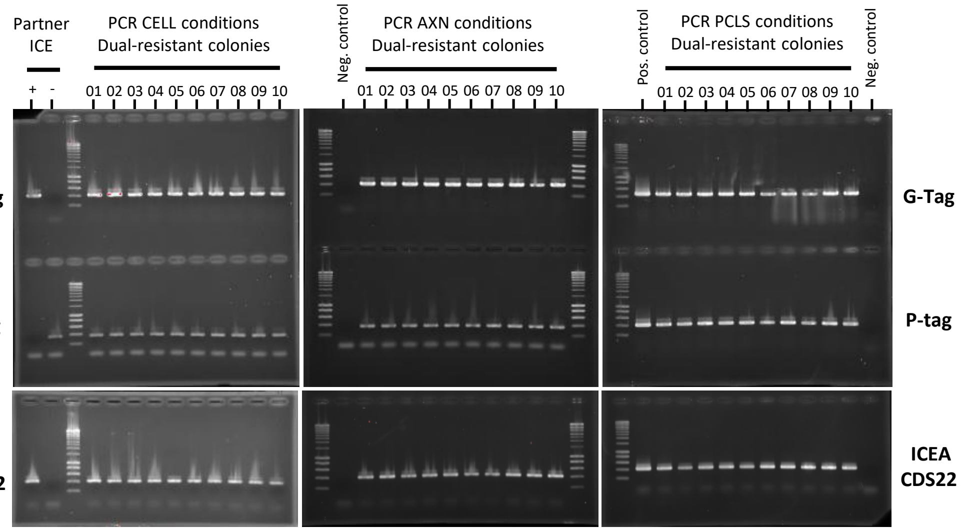


**Mating E** (5632T x PG2G)

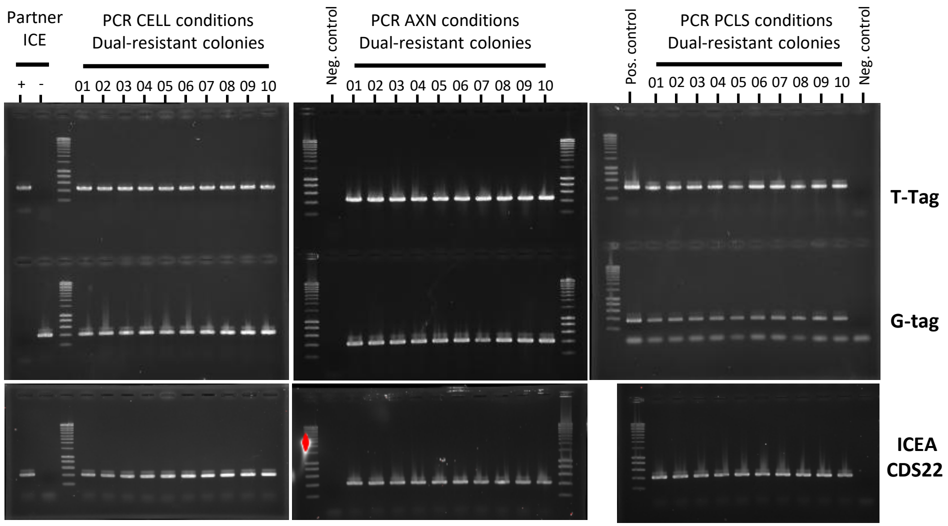


**Mating F** (J228[ICEB]G x pRM16P)

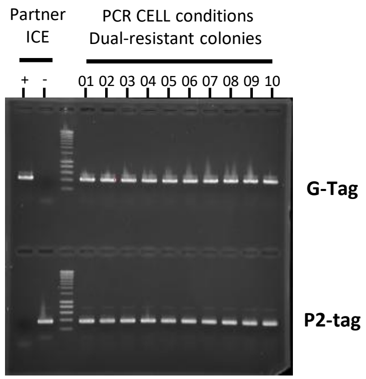


**Figure S1. PCR characterization of dual-resistant colonies.**
PCR amplification of 10 dual-resistant colonies selected from mating A, B, E and F (Fig. 1) under axenic (AXN), cell (CELL) and organotypic (PCLS) mating conditions. PCR targets are the G-tag, P-tag, P2-tag, T-tag and ICEA CDS22 (Table S1). ICE partners were used as controls.
